# Traumatic Brain Injury-Associated Micrgoglia Adopt Longitudinal Transcriptional Changes Consistent with Long-Term Depression of Synaptic Strength

**DOI:** 10.1101/422071

**Authors:** Hadijat M. Makinde, Talia B. Just, Deborah R. Winter, Steven J. Schwulst

## Abstract

Traumatic brain injury (TBI) is an under-recognized public health threat. Even mild brain injury, or concussions, may lead to long-term neurologic impairment. Microglia play a fundamental role in the development and progression of this subsequent neurologic impairment. Despite this, a microglia-specific injury signature has yet to be identified. In the current study we hypothesized that TBI-associated microglia would adopt longitudinal changes in their transcriptional profile associated with pathways linked to the development of motor, cognitive, and behavioral disorders. C57BL/6 mice underwent TBI via a controlled cortical impact and were followed longitudinally. FACSorted microglia from TBI mice were subjected to RNA-sequencing at 7, 30, and 90 days post-injury. We identified 4 major patterns of gene expression corresponding to the host defense response, synaptic potentiation, lipid remodeling, and membrane polarization. In particular, significant upregulation of genes involved in long-term synaptic potentiation including Ptpn5, Shank3, and Sqstm1 were observed offering new insight into a previously unknown role of microglia in the weakening of synaptic efficacy between neurons after brain injury.

## Introduction

Traumatic brain injury is a growing and under recognized public health threat. The CDC estimates nearly 2 million people sustain a traumatic brain injury (TBI) each year in the United States, contributing to over 30% of all injury related deaths ^1,2^. In fact, TBI related healthcare expenditures near 80 billion dollars annually with an average cost of 4 million dollars per person surviving a severe TBI ^3-5^. The impact of TBI is highlighted not only by its high mortality rate but also by the significant long-term complications suffered by its survivors with the progressive development of motor, cognitive, and behavioral disorders ^6-10^. Even subconcussive events, those resulting in subclinical brain dysfunction without the typical symptoms of concussion, may lead to long-term neurologic impairment ^11,12^. The immune response to TBI plays a fundamental role the development and progression of subsequent neurodegenerative disease and represents a complex interplay between the injured brain and the resident immune cells of the brain — the microglia ^13^. The current manuscript is focused on developing a cell-type-specific understanding of the microglial response to injury over time. Here we highlight the major trends in gene expression in response to TBI.

TBI triggers a robust pro-inflammatory response within the injured brain. The degree of this initial pro-inflammatory response has significant value in predicting more long-term outcomes after TBI ^14-16^. Even after the acute inflammatory response has resolved, several studies demonstrated residual long-lasting inflammation within the brain in both animal models as well as in patients ^17,18^. One of the main drivers of this continued inflammation is the persistence of activated microglia—characterized by thickening and retraction of their ramified processes, increased IL-1 and IL-6 with concomitant decreases in IL-4 and IL-10, and increased expression of pro-oxidant genes with a reduction of growth and antioxidant genes. A recent study showed an increased inflammatory profile of microglia that persisted for up to 12 months after injury in a murine model of TBI ^19^. Furthermore, this continued inflammation is associated with lesion volume expansion and loss of neurons in the hippocampus ^18-22^. Even once the acute inflammatory process has resolved and infiltrated monocyte-derived macrophages are no longer present within the injured brain, microglia have the potential to remain activated for years after the initial insult ^19,23^. A functional consequence of this constitutive activation after brain trauma is an exaggerated neuroinflammatory response to otherwise benign secondary stimuli such a subsequent subclinical head injury ^10,24-27^. This may be the mechanism by which patients who have sustained a concussion are more susceptible to subsequent concussions ^28-30^. Nonetheless, the molecular mechanisms resulting in the constitutive activation of microglia remain elusive ^21^. Therefore, in the current study, we aimed to study the transcriptional dynamics of constitutively activated microglia over the course of brain injury.

The first step towards any cell-specific transcriptional analysis relies on obtaining sufficient cells of interest with the highest purity. The historical standard for distinguishing between microglia and infiltrating macrophages is immunohistochemistry. Although immunohistochemistry is useful for assessing morphology, proliferation, and sites of activation it has a number of drawbacks limiting its use ^31^. Several investigative groups have focused on this problem including column free magnetic separation and CD11b immunomagnetic enrichment combined with the differential expression of CD45 with flow cytometry ^31-33^. However, CD45 expression has been reported to vary depending on the pathologic condition; thus, reliable separation of microglia from peripheral myeloid cells is impaired ^34,35^. To overcome this issue, fluorescently marked myeloid cells, such as CX3CR1^+/GFP^/CCR2^+/RFP^, have been used. However, these mice are on mixed backgrounds, which could greatly affect the immune response. Furthermore, the presence of contaminated nonclassical monocytes could not be excluded ^36,37^. Therefore, we use head-shielded bone marrow chimeric mice with CD45.1 cells in the circulation and CD45.2 microglia in the brain allowing definitive and unambiguous differentiation between microglia and infiltrating bone-marrow derived myeloid cells as previously described by our laboratory ^38^. To the best of our knowledge, no cell-type-specific study has been conducted to specifically identify transcriptional changes in isolated populations of microglia over the course of TBI. In the current study we demonstrate that TBI-associated microglia adopt longitudinal changes in their transcriptional profile associated with pathways linked to the development of motor, cognitive, and behavioral disorders.

## Results

### Global patterns of gene expression from isolated populations of microglia over the time course of TBI

The neuroinflammatory response to TBI is central to both neuroprotection and neurotoxicity after injury, but attempts to broadly target immune activation have been unsuccessful in improving outcomes in TBI patients ^39-42^. Because of this, there has been a growing interest in investigating the microglial transcriptome after brain injury. Attempts thus far have been plagued by the use of whole brain homogenates rather than individual cell types as well as the use of microarray analyses restricted to a limited number of genes in the inflammatory response ^43,44^. Therefore, we combined our ability to discriminate and sort microglia from infiltrating monocytes and macrophages with unbiased transcriptional profiling (RNA-seq) on FACSorted microglia from 7, 30, and 90 days after TBI. We defined 396 genes that change expression across the time course (see methods). We clustered these differentially expressed genes into 4 main patterns of expression (**Fig. 1)**. Clusters 2 represents genes involved in synaptic plasticity and is progressively upregulated over the course of injury. Cluster 4 represents genes involved in the regulation of membrane polarization with gene expression that is upregulated at 30 days post-injury and then downregulated by 90 days post injury. Clusters 1 and 3 represent genes involved in the host response to injury and lipid remodeling and both are progressively downregulated over the course of injury. These data provide new insights into the biology of microglial activation over the course of TBI.

**Figure 1.**
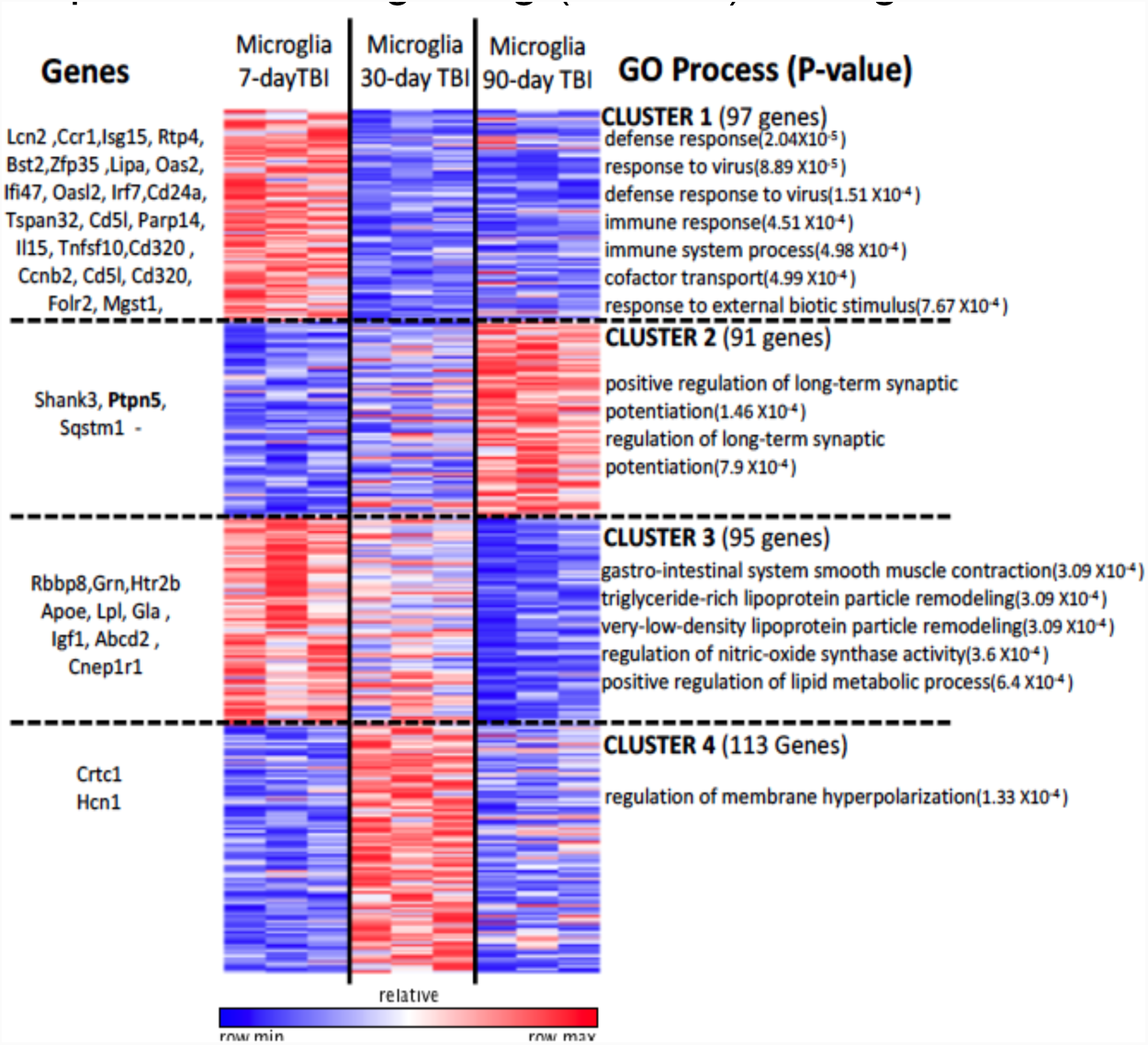
Microglia from TBI mice show distinct time-dependent transcriptional profiles. K-means clustering: Four clusters representing 392 genes are shown with distinct time specific expression patterns. Averaged log_2_(CPM+1) ≥3 of genes in each group that had a log foldchange of at least 2 was used to generate the heat map. GO processes associated with the genes in each cluster was determined using Gorilla database.

### Pairwise comparison of microglia gene expression between time points post-TBI

To further evaluate the gene expression over time in TBI-associated microglia, we investigated differentially expressed genes between microglia across time points (**Fig. 2**). We found 187 genes between days 7 and 30 that are positively differentiated and are likely involved in channel transport activity, as well as defense response. Several genes including genes involved in immune response and tissue repair such as Tnfsf10, Bst2, Igf1, and Ccr1 were upregulated at the earlier time points. However, genes implicated in neurodegenerative diseases such as Ptpn5, Shank3, and Sqstm1 were upregulated by 90 days compared to 7 days post TBI. This expression profile is indicative of the chronic inflammatory environment in the TBI brain following injury. To determine whether gene expression trends at 30 days were predictive of 90 days post-TBI, we compared the fold changes between 7 vs. 30 days post-TBI with those between 7 vs. 90 days post-TBI (**Fig. 3**). We find that these gene expression change were significantly correlated (R=-0.537, p=2.2 × 10^-16^). This indicates that genes which were upregulated at 90 days post-TBI had already started to increase as early as 30 days post-injury. Conversely, those genes that were downregulated at 30 days post-TBI remained downregulated or continued to fall over time.

**Figure 2.**
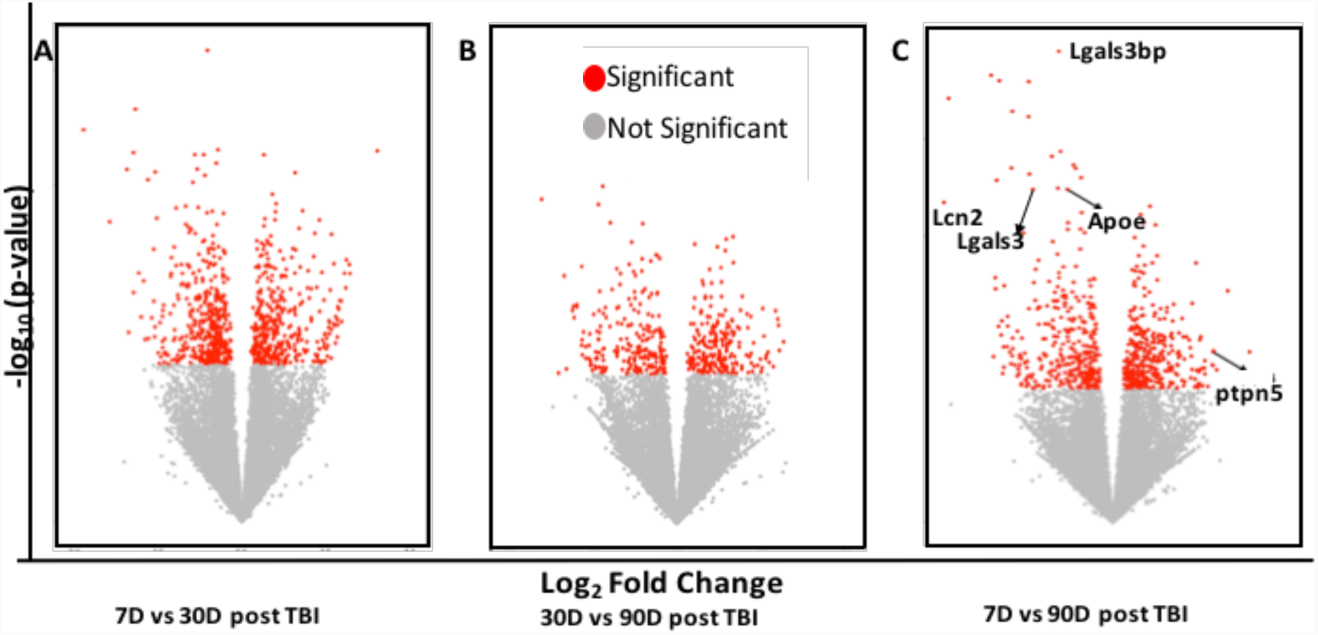
Differentially expressed genes over the time course of injury. Volcano plots of differentially expressed genes in microglia between 7D and 30D post-TBI (A), 30D and 90D post-TBI (B), and 7D and 90D post-TBI (C). N=3 in each group. Genes with a p-value <0.05 are shown in red.

**Figure 3.**
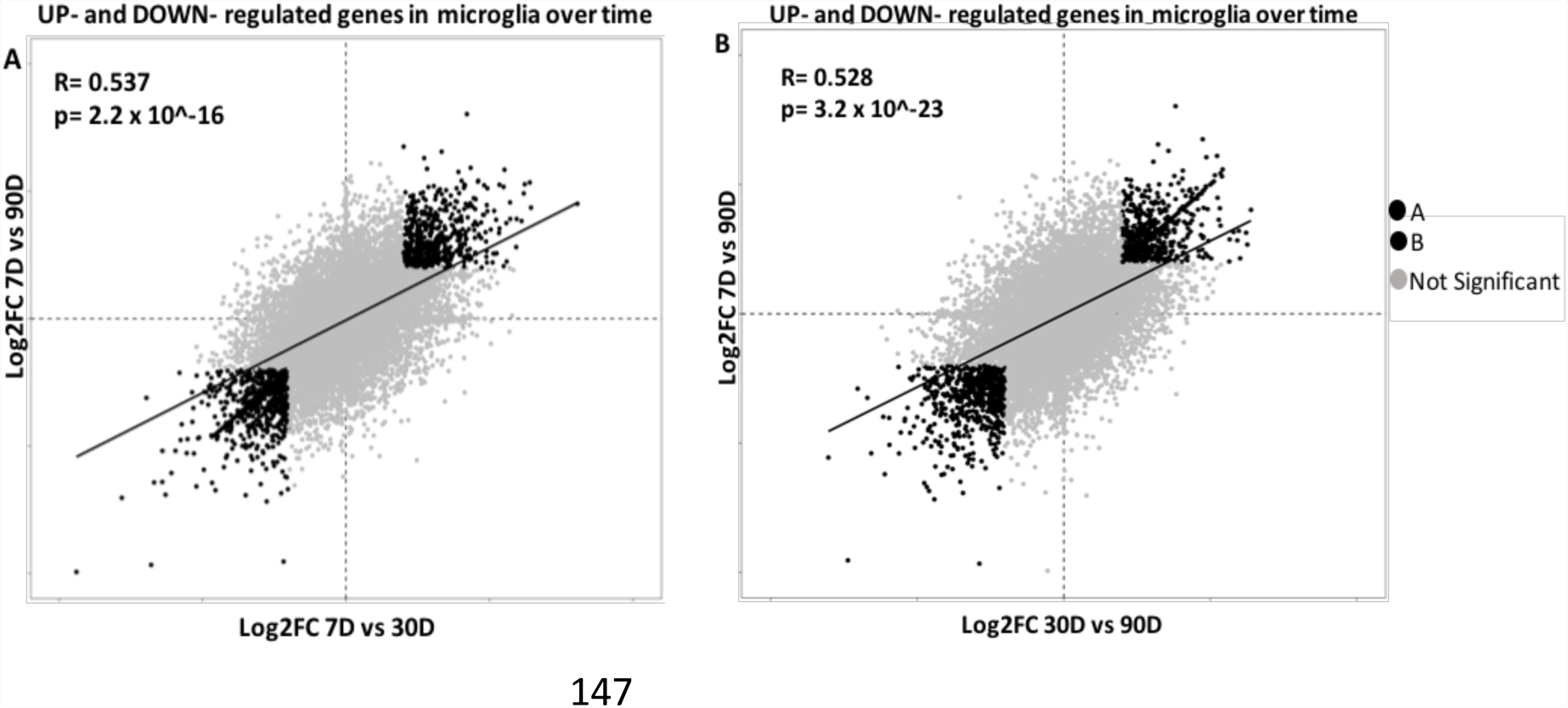
Scatter plot of fold change in CPM of microglia over time after TBI. Scatter plot to examine the relationship between the genes that are up-regulated or down-regulated in the microglia of mice at A) 7 vs 30 days post-TBI compared to 7 vs 90 days post-TBI and at B) 30 vs 90 days post-TBI compared to 7 vs 90 days. A log2 fold change of 1 is equal to a 2-fold change.

### Trem2-APOE pathway is not upregulated over the course of traumatic brain injury

Microglia are essential to brain homeostasis but lose this homeostatic function in a number of neurodegenerative disease processes. There has been considerable interest in the Trem2-APOE pathway in the generation of a neurodegenerative microglial phenotype in both Alzheimer’s Disease (AD) and multiple sclerosis (MS). In fact, recent data has identified the Trem2-APOE pathway as a pivotal regulator of microglial phenotype in both of these disease processes ^45^. Therefore, we aimed to determine if the Trem2-APOE pathway was a major regulator of microglial phenotype after TBI. However, unlike other neurodegenerative processes, our data demonstrates no significant change in TREM2 expression as well as a progressive decrease in APOE expression over the course of TBI (**Fig. 4**). These seemingly contradictory results emphasize the need for microglia-specific transcriptional studies in the setting TBI.

**Figure 4.**
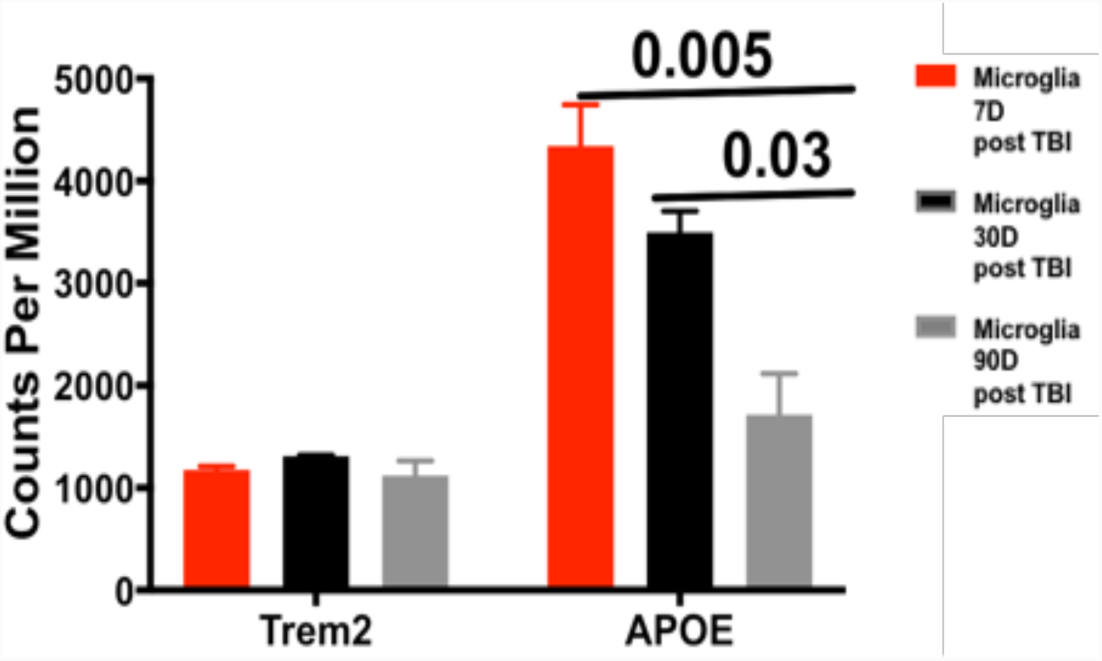
Microglial Trem2 and APOE expression over the course of TBI. Microglial Trem2 did not significantly change over the course of brain injury. APOE expression progressively decreased over time following TBI. Ordinary one way ANOVA with Bonferroni’s multiple comparison test.

### Microglial STEP expression is progressively upregulated over the course of brain injury

While our data failed to demonstrate a common pathway with Alzheimer’s disease and multiple sclerosis through Trem2-APOE, a closer examination of Cluster 2 (**Fig. 1**) revealed a number of upregulated genes associated with long-term synaptic potentiation, including PTPN5 also known as STEP (STriatial-Enriched protein tyrosine Phosphatase). STEP is a brain-specific phosphatase and is highly expressed within the cortex, hippocampus, and amygdala ^46,47^. Our data show that expression of STEP is progressively increased in microglia over time following TBI (**Fig. 5**). STEP is critical in the long-term depression, or weakening, of synaptic efficacy between neurons—a process fundamental to learning, memory, and cognition ^48,49^. When STEP activity is elevated, several substrates are inactivated resulting in the internalization of NMDA/AMPA glutamate receptors ^50^. This disrupts synaptic function and contributes to cognitive deficits ^49,51^. In fact, elevated STEP is associated with the pathophysiology of Alzheimer’s disease, schizophrenia, and ischemic brain injury in both human cortex and mouse models ^52-57^. These data suggest that STEP may be one of the common molecular pathways connecting TBI with other known neurodegenerative disorders.

**Figure 5.**
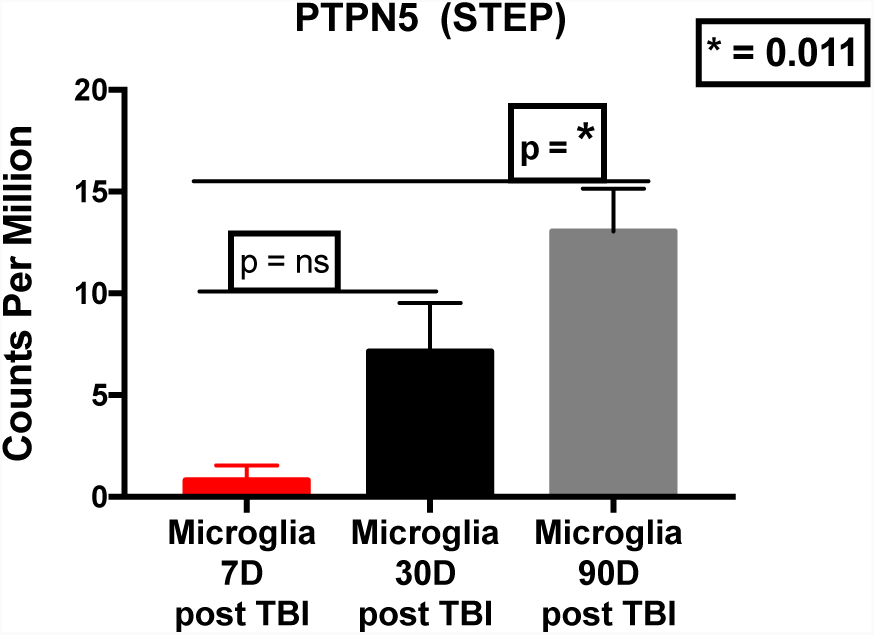
Microglial STEP expression is progressively upregulated over the course of brain injury. STEP expression from FACSorted microglia progressively increased from 7 days post-TBI to 90 days post-TBI; *p*=0.01, Ordinary one way ANOVA with Bonferroni’s multiple comparison test.

## Discussion

Microglia are the resident innate immune cells of the CNS. They are ontologically distinct from peripheral bone marrow-derived monocytes and macrophages, arising from the yolk sac as opposed to the developing liver in the embryo ^58^. In fact, microglia rely on a distinctive set of transcription factors during development resulting in a lineage of tissue macrophages (microglia) derived from the yolk sac that are genetically distinct from bone marrow-derived macrophages ^37,59-62^. Additionally, microglia are self-renewing suggesting that monocyte-derived macrophages do not contribute to the maintenance of the mature microglia pool ^63,64^. Distinct developmental origin and renewal mechanisms may suggest that microglia possess discrete functions in pathological processes ^58^. Despite this, the cellular mechanisms by which microglia promote or attenuate the progression of injury are largely unknown ^65^. Our Data are the first we know of to use unbiased transcriptional profiling of isolated populations of microglia to define the genes/pathways/signatures involved in the generation of TBI-associated microglia over the course of injury.

To adequately capture the heterogeneity and complexity of microglia at different stages of injury, a comprehensive, genome-wide sampling of individual cell types is required ^37,66^. A cell-specific delineation of innate immune function based on transcriptional profiling in TBI has yet to be undertaken. Therefore, we combined our ability to discriminate and sort microglia from infiltrating monocytes and monocyte-derived macrophages with unbiased transcriptional profiling (RNA-seq) on FACSorted microglia. Our analysis identified 4 sequentially upregulated or downregulated gene clusters involved in processes such as the host defense response, synaptic potentiation, lipid remodeling, and membrane polarization each providing new insights into the biology of microglial activation over the course of TBI (**Fig. 1**). While there has recently been considerable recent interest in the Trem2-APOE pathway in the generation of a neurodegenerative microglial phenotype in Alzheimer’s Disease and multiple sclerosis ^45^, our data demonstrates no significant change in Trem2 expression as well as a progressive decrease in APOE expression over the course of TBI (**Fig. 4**). However, an examination of cluster II revealed a number of genes associated with long-term synaptic potentiation, including PTPN5 also known as STEP (STriatial-Enriched protein tyrosine Phosphatase). STEP is a brain-specific phosphatase that is highly expressed within the striatum, cortex, hippocampus, and amygdala ^46,47^. STEP is critical in the long-term depression, or weakening, of synaptic efficacy between neurons—a process fundamental to learning, memory, and cognition ^48,49^. Elevated STEP is associated with the pathophysiology of Alzheimer’s disease (AD), schizophrenia, and ischemic brain injury in both human cortex and mouse models ^52-57^. In fact, genetic knockout of STEP reverses many of the cognitive and behavioral deficits in AD models ^53,67^. To the best of our knowledge, STEP expression has never been studied within the context of TBI. Our RNA-seq analysis demonstrates a 13-fold increase in microglial expression of STEP over the course of injury (**Fig. 5**). Previous studies have shown that STEP affects neuronal communication by opposing synaptic strengthening. High levels of STEP are believed to disrupt synaptic function and to contribute to learning deficits in neurodegenerative disease ^52,68^. When STEP activity is elevated, several substrates are inactivated resulting in the internalization of NMDA/AMPA glutamate receptors ^50^. This disrupts synaptic function and contributes to cognitive deficits ^49,51^. In other words, STEP activation modulates learning and memory by removing glutamate receptors from synaptic membranes. This important discovery suggests that TBI may share a common molecular pathway with several other cognitive disorders previously regarded as distinct. These data are remarkable in demonstrating the power of longitudinal transcriptional profiling, which provides important biologic insights into the state of microglial processes even in the complex and dynamic model of traumatic brain injury. Furthermore, these data strongly implicate longitudinal changes in microglial gene expression in the development of long-term neurocognitive changes.

In conclusion, our data demonstrate that TBI-associated microglia adopt longitudinal transcriptional changes consistent with long-term depression of synaptic strength. The contribution of altered microglial gene expression to the pathogenesis of TBI has not been previously investigated. Our data suggest that TBI-associated microglia may play a previously unknown role in the weakening of synaptic efficacy between neurons after brain injury. As a result, learning, memory, and cognitive performance may all be affected leading to the resultant long-term neurocognitive impairments seen after TBI. Moving forward it will be important to study larger cohorts of brain-injured mice during both the acute and chronic phase of TBI. Furthermore, it has been shown that microglia display different transcriptional identities depending on the brain region in which they reside as well as their age ^69^. This will necessitate side-by-side comparison with age-matched naïve control mice to account for transcriptional changes associated with aging. Additionally, a single-cell RNA-seq approach may be required to account for inherent microglial heterogeneity at the site of injury. This could allow for the identification of novel microglial subpopulations within and surrounding the site of injury. Regardless of the techniques used, once the molecular mechanisms underlying the transcriptional changes in microglia after injury are further delineated, targeting the microglial response after TBI may soon represent a target for future therapeutic intervention.

## Methods

### Mice

All procedures were approved by the Northwestern University Institutional Animal Care and Use committee and all experiments were carried out in accordance with the ARRIVE guidelines on the reporting of in vivo experiments. Two mouse strains were used, they are C57BL/6 and B6.SJL-*Ptprc^a^Pepc^b^*/BoyJ (CD45.1). All mice were purchased from the Jackson Laboratory and housed at a barrier facility at the Center for Comparative Medicine at Northwestern University (Chicago, IL, USA). Sixteen-week-old mice were used for all experiments.

### Shielded Bone Marrow Chimeras

Bone marrow was aseptically harvested from tibias and femurs, from 8-week-old B6.CD45.1 donor mice, erythrocytes were lysed and the cells were counted using a Countess automated cell counter. 8 weeks old B6.*CD45.2* mice received a single 1000-cGy γ-irradiation dose using a Cs-137-based Gammacell 40 irradiator. The mice heads were shielded with a lead bar so as to deliver the irradiation to the body only. 6 hours after shielded irradiation, busulfan (30 mg/kg) was administered to completely ablate the bone marrow of the recipient mice. Donor bone marrow (CD45.1) was transplanted 12 hours after busulfan ablation. Shielded bone marrow chimeras were maintained on antibiotics trimethoprim/sulfamethoxazole (40 mg/5 mg, respectively). Eight weeks after irradiation, 95% of the circulating monocytes were of donor origin (**Fig. 6**) ^38^.

**Figure 6.**
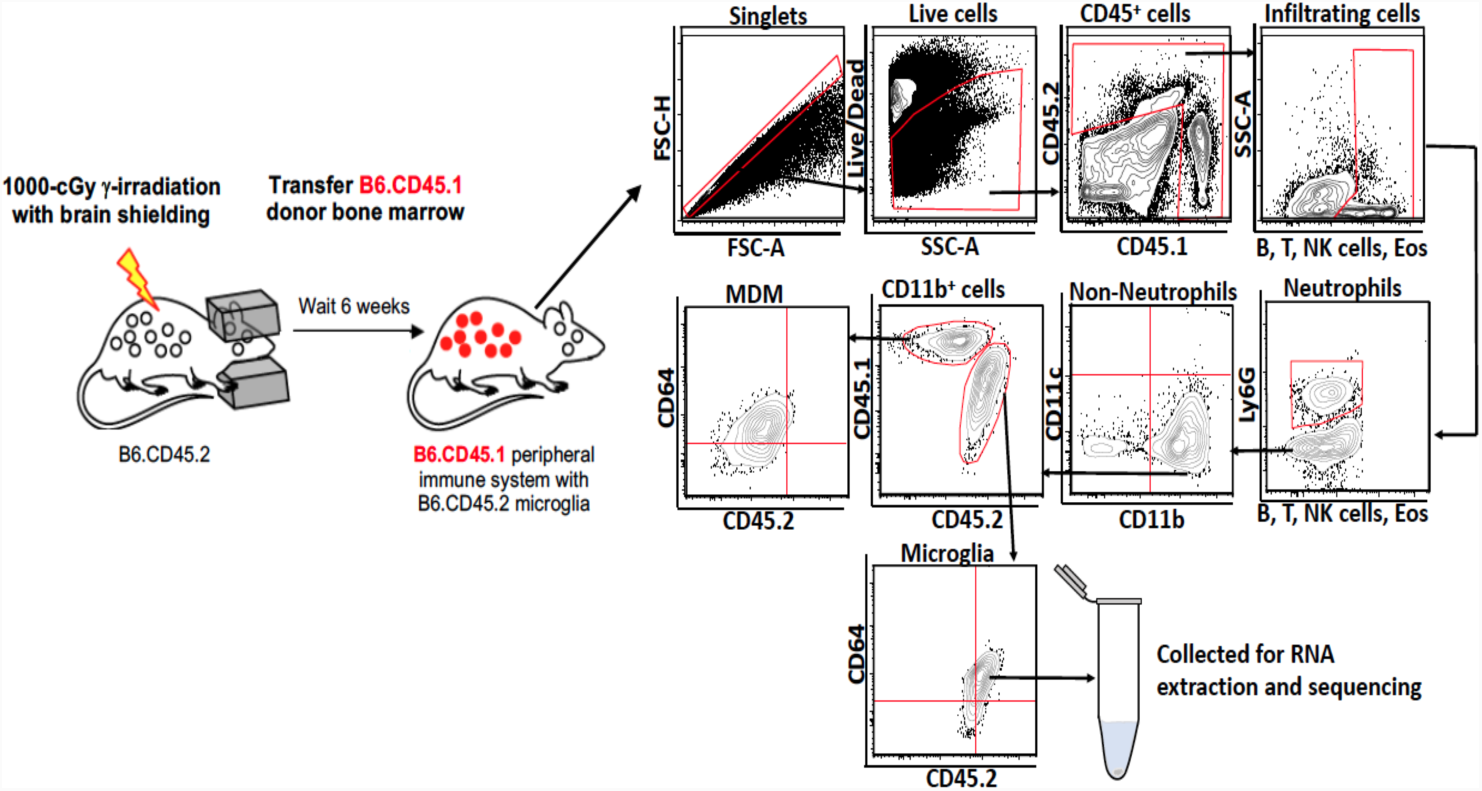
Microglia from head-shielded bone marrow chimeric mice are host origin. Brains isolated from chimeric mice post TBI were analyzed by flow cytometry. CD45.1^neg^‘B, T, NK cells, Eosinophils’ acted as the gating control for CD45.2^hi^, showing that resident microglia (CD45^lo^ can be unambiguously differentiated from infiltrating monocyte-derived macrophages (CD45.1^+^). Arrows indicated the directionality of gating.

### Controlled cortical impact

Controlled cortical impact was induced as previously described by our laboratory ^38^. In brief, mice were anesthetized with 100 mg/kg Ketamine and 10 mg/kg Xylazine via intraperitoneal injection. A 1cm scalp incision was performed and a 5mm craniectomy was performed 2 mm left of the sagittal suture and 2 mm rostral to the coronal suture. The dura was left intact. Mice were then placed in a stereotaxic operating frame and the impactor (Leica Biosystems Inc., Buffalo Grove, IL) was maneuvered into position. A controlled cortical impact was then applied with a 3mm impacting tip at a velocity of 2.5m/s and an impacting depth of 2mm with the dwell time set at 0.1s. Immediately following injury all animals had their scalps sealed with VetBond (3M). All animals received post injury analgesia with Buprenorphine SR via subcutaneous injection and were allowed to recover in separate cages over a warming pad. Mice were euthanized at 7, 30, and 90 days post TBI and brains were harvested.

### Tissue preparation and fluorescence activated cell sorting

Immediately following euthanasia, mice were transcardially perfused first with ice cold PBS. The brains were then excised and place in ice cold HBSS until time to process. The brains were weighed, cut into pieces and placed into C-tubes containing digestion buffer (2.5 mg/mL Liberase TL (Roche, Basel, Switzerland), and 1 mg/mL of DNase I in HBSS). The C-tubes were placed on a MACS dissociator (Miltenyi Biotec) and run on the M_Brain_3 protocol, after which they were placed in an incubator for 30 minutes at 37°C with shaking at 200 rpm. After incubation, the C-tubes were placed back on the MACS dissociator and run on the same protocol as before. The cells released were then passed through a 40 μm nylon mesh and washed with 100ml of wash buffer (1% BSA in HBSS) per brain sample. Microglia and infiltrating cells were isolated using a 30/70 percoll gradient (Percoll Plus, GE Healthcare). The cells collected from the interphase of the gradient were washed with HBSS and counted using Countess automated cell counter (Invitrogen); dead cells were discriminated using trypan blue. Cells were stained with live/dead Aqua (Invitrogen) viability dye, incubated with Fc-Block (BD Bioscience) and stained with fluorochrome-conjugated antibodies (**Table 1**). Data were acquired on a BD FACSAria cell sorter (BD Biosciences, San Jose, CA), and microglia were sorted for further analyses. “Fluorescence minus one” controls were used when necessary to set up gates. Pelleted sorted cells were immediately lysed in extraction buffer from a PicoPure RNA isolation kit (Arcturus Bioscience), and lysates were stored at −80°C until RNA was extracted. Analysis of the flow cytometric data was performed using Flowjo software (TreeStar, Ashland, OR).

**Table 1.** List of antibody conjugated fluorochromes used to differentiate microglia from infiltrating leukocytes.

| Antibody | Fluorochrome | Company and Clone |
| --- | --- | --- |
| CD45.1 | FITC | A-20 / BD Bioscience |
| CD45.2 | BV421 | 104 / Biolegend |
| CD64 | APC | X54-5/7.1/ BD Bioscience |
| CD11b | APC- Cy7 | M1/70 / BD Biosciences |
| CD11c | PE-Cy7 | HL3 / BD Bioscience |
| Ly6G | Alexa Flour 700 | 1A8 / BD Bioscience |
| MHC II | Percp- Cy 5.5 | M5/114.15.2 / Biolegend |
| Siglec H | PE | 551 / Biolegend |
| B220 | PECF594 | RA3-6B2 / BD Bioscience |
| Siglec F | PECF594 | E50-2440 / BD Bioscience |
| CD4 | PECF594 | RM4-5/ BD Bioscience |
| CD8 | PECF594 | 53-6.7 / BD Bioscience |
| NK1.1 | PECF594 | PK136/ BD Bioscience |
| Viability | e-bioscience Fixable Viability Dye eFluor 506 | Invitrogen |

### RNA sequencing

RNA from the FACSorted microglia of brain-injured mice were extracted using a PicoPure RNA isolation kit according to manufacturer’s instructions. Sample quality control, processing, and library preparation were performed by the Northwestern University Next Generation Sequencing Core (NUSeq). RNA quality and quantity were measured using Agilent High Sensitivity RNA ScreenTape System (Agilent Technologies). RNA sequencing (RNA-seq) libraries were prepared from 3ng of total RNA using the QuantSeq 3’ biased mRNA-seq Library Prep Kit for Illumina (Lexogen). DNA libraries were sequenced on an Illumina NextSeq 500 instrument with a target read depth of ~20 million reads per sample.

### RNA-seq analysis

Raw sequencing files were first de-multiplexed using bcl2fastq. The resulting fastq files were trimmed of low-quality reads and bases, polyA tails, and adaptors using bbduk (http://jgi.doe.gov/data-and-tools/bb-tools/). The trimmed fastq files were aligned to the mouse reference genome (mm10, Genome Reference Consortium GRCm38) using the STAR (Spliced Transcripts Alignment to a Reference) algorithm ^70^. HTSeq was run on the resulting BAM files to provide raw gene counts. Raw gene counts for each sample were merged into a single gene expression table and normalized for read depth using counts per million (CPM). The three highest quality samples, based on RNA quality and library quality from each experimental group were included for subsequent analyses. For the RNA-seq analysis, we focused on the expressed genes which were defined as average log CPM (log_2_(CPM+1)) expression > 4 in each experimental group. For visualization, GENE-E (https://software.broadinstitute.org/GENE-E/) was used to perform K-means clustering (K=4) on differentially expressed genes across all time points as defined by ANOVA test (p<0.05) across any two groups shown in the heatmap. Gene Ontology associations and the related p-values were determined by GO analysis (by GOrilla—[Gene Ontology enRIchment *anaLysis* and visuaLizAtion tool]).

^71^. Pairwise differential genes between time points were determined using DEseq2. Volcano plots were generated using the log2 fold change of normalized gene counts between microglia at different time points on the x-axis and corresponding p-values (-log10) from DEseq2 on the y-axis. Plots were generated using the ggplot2 package from R Studio software.

## Data Availability

The data that support the findings of this study are available from the corresponding author on request. RNA sequencing data is available through the NCBI Sequence Read Archive (SRA accession number: SRP160379).

## Author Contributions

S.S and H.M wrote the main manuscript text and prepared all figures. H.M, T.J and S.S carried out the experiments. H.M and D.W performed the computational analysis of the sequencing data.

## References

1. Faul, M. Traumatic Brain Injury in the United States: Emergency Department Visits, Hospitalizations and Deaths 2002-2006. (Centers for Disease Control and Prevention, National Center for Injury Prevention and Control, Atlanta (GA), 2010).

2. Roozenbeek, B., Maas, A. I. & Menon, D. K. Changing patterns in the epidemiology of traumatic brain injury. Nat Rev Neurol 9, 231–236, doi:10.1038/nrneurol.2013.22 (2013).

3. Corso, P., Finkelstein, E., Miller, T., Fiebelkorn, I. & Zaloshnja, E. Incidence and lifetime costs of injuries in the United States. Injury prevention : journal of the International Society for Child and Adolescent Injury Prevention 12, 212–218, doi:10.1136/ip.2005.010983 (2006).

4. Pearson, W. S., Sugerman, D. E., McGuire, L. C. & Coronado, V. G. Emergency department visits for traumatic brain injury in older adults in the United States: 2006-08. The western journal of emergency medicine 13, 289–293, doi:10.5811/westjem.2012.3.11559 (2012).

5. Whitlock, J. A., Jr. & Hamilton, B. B. Functional outcome after rehabilitation for severe traumatic brain injury. Archives of physical medicine and rehabilitation 76, 1103–1112 (1995).

6. Schwarzbold, M. et al. Psychiatric disorders and traumatic brain injury. Neuropsychiatric disease and treatment 4, 797–816 (2008).

7. Whelan-Goodinson, R., Ponsford, J., Johnston, L. & Grant, F. Psychiatric disorders following traumatic brain injury: their nature and frequency. The Journal of head trauma rehabilitation 24, 324–332, doi:10.1097/HTR.0b013e3181a712aa (2009).

8. Peskind, E. R., Brody, D., Cernak, I., McKee, A. & Ruff, R. L. Military- and sports-related mild traumatic brain injury: clinical presentation, management, and long-term consequences. The Journal of clinical psychiatry 74, 180-188; quiz 188, doi:10.4088/JCP.12011co1c (2013).

9. Martin, L. A., Neighbors, H. W. & Griffith, D. M. The experience of symptoms of depression in men vs women: analysis of the National Comorbidity Survey Replication. JAMA psychiatry 70, 1100–1106, doi:10.1001/jamapsychiatry.2013.1985 (2013).

10. Makinde, H. M., Just, T. B., Cuda, C. M., Perlman, H. & Schwulst, S. J. The Role of Microglia in the Etiology and Evolution of Chronic Traumatic Encephalopathy. Shock 48, 276–283, doi:10.1097/SHK.0000000000000859 (2017).

11. Belanger, H. G., Vanderploeg, R. D. & McAllister, T. Subconcussive Blows to the Head: A Formative Review of Short-term Clinical Outcomes. J Head Trauma Rehabil 31, 159–166, doi:10.1097/HTR.0000000000000138 (2016).

12. Carman, A. J. et al. Expert consensus document: Mind the gaps-advancing research into short-term and long-term neuropsychological outcomes of youth sports-related concussions. Nat Rev Neurol 11, 230–244, doi:10.1038/nrneurol.2015.30 (2015).

13. Hernandez-Ontiveros, D. G. et al. Microglia activation as a biomarker for traumatic brain injury. Front Neurol 4, 30, doi:10.3389/fneur.2013.00030 (2013).

14. Kumar, R. G. et al. Acute CSF interleukin-6 trajectories after TBI: associations with neuroinflammation, polytrauma, and outcome. Brain Behav Immun 45, 253–262, doi:10.1016/j.bbi.2014.12.021 (2015).

15. Winter, C. D., Pringle, A. K., Clough, G. F. & Church, M. K. Raised parenchymal interleukin-6 levels correlate with improved outcome after traumatic brain injury. Brain 127, 315–320, doi:10.1093/brain/awh039 (2004).

16. Thelin, E. P. et al. Monitoring the Neuroinflammatory Response Following Acute Brain Injury. Front Neurol 8, 351, doi:10.3389/fneur.2017.00351 (2017).

17. Kumar, R. G., Boles, J. A. & Wagner, A. K. Chronic Inflammation After Severe Traumatic Brain Injury: Characterization and Associations With Outcome at 6 and 12 Months Postinjury. The Journal of head trauma rehabilitation 30, 369–381, doi:10.1097/HTR.0000000000000067 (2015).

18. Smith, D. H., Johnson, V. E. & Stewart, W. Chronic neuropathologies of single and repetitive TBI: substrates of dementia? Nature reviews. Neurology 9, 211–221, doi:10.1038/nrneurol.2013.29 (2013).

19. Loane, D. J., Kumar, A., Stoica, B. A., Cabatbat, R. & Faden, A. I. Progressive neurodegeneration after experimental brain trauma: association with chronic microglial activation. J Neuropathol Exp Neurol 73, 14–29, doi:10.1097/NEN.0000000000000021 (2014).

20. Lozano, D. et al. Neuroinflammatory responses to traumatic brain injury: etiology, clinical consequences, and therapeutic opportunities. Neuropsychiatric disease and treatment 11, 97–106, doi:10.2147/NDT.S65815 (2015).

21. Perry, V. H. & Holmes, C. Microglial priming in neurodegenerative disease. Nat Rev Neurol 10, 217–224, doi:10.1038/nrneurol.2014.38 (2014).

22. Henry, C. J., Huang, Y., Wynne, A. M. & Godbout, J. P. Peripheral lipopolysaccharide (LPS) challenge promotes microglial hyperactivity in aged mice that is associated with exaggerated induction of both pro-inflammatory IL-1beta and anti-inflammatory IL-10 cytokines. Brain Behav Immun 23, 309–317, doi:10.1016/j.bbi.2008.09.002 (2009).

23. Johnson, V. E. et al. Inflammation and white matter degeneration persist for years after a single traumatic brain injury. Brain 136, 28–42, doi:10.1093/brain/aws322 (2013).

24. Field, R., Campion, S., Warren, C., Murray, C. & Cunningham, C. Systemic challenge with the TLR3 agonist poly I:C induces amplified IFNalpha/beta and IL-1beta responses in the diseased brain and exacerbates chronic neurodegeneration. Brain Behav Immun 24, 996–1007, doi:10.1016/j.bbi.2010.04.004 (2010).

25. Ohmoto, Y. et al. Variation in the immune response to adenoviral vectors in the brain: influence of mouse strain, environmental conditions and priming. Gene Ther 6, 471–481, doi:10.1038/sj.gt.3300851 (1999).

26. McColl, B. W., Rothwell, N. J. & Allan, S. M. Systemic inflammatory stimulus potentiates the acute phase and CXC chemokine responses to experimental stroke and exacerbates brain damage via interleukin-1- and neutrophil-dependent mechanisms. The Journal of neuroscience : the official journal of the Society for Neuroscience 27, 4403–4412, doi:10.1523/JNEUROSCI.5376-06.2007 (2007).

27. Schroder, K., Sweet, M. J. & Hume, D. A. Signal integration between IFNgamma and TLR signalling pathways in macrophages. Immunobiology 211, 511–524, doi:10.1016/j.imbio.2006.05.007 (2006).

28. Muccigrosso, M. M. et al. Cognitive deficits develop 1month after diffuse brain injury and are exaggerated by microglia-associated reactivity to peripheral immune challenge. Brain Behav Immun 54, 95–109, doi:10.1016/j.bbi.2016.01.009 (2016).

29. Weil, Z. M., Gaier, K. R. & Karelina, K. Injury timing alters metabolic, inflammatory and functional outcomes following repeated mild traumatic brain injury. Neurobiol Dis 70, 108–116, doi:10.1016/j.nbd.2014.06.016 (2014).

30. Foris, L. A. & Donnally, I. C. in StatPearls (2017).

31. Nikodemova, M. & Watters, J. J. Efficient isolation of live microglia with preserved phenotypes from adult mouse brain. Journal of neuroinflammation 9, 147, doi:10.1186/1742-2094-9-147 (2012).

32. Bedi, S. S., Smith, P., Hetz, R. A., Xue, H. & Cox, C. S. Immunomagnetic enrichment and flow cytometric characterization of mouse microglia. Journal of neuroscience methods 219, 176–182, doi:10.1016/j.jneumeth.2013.07.017 (2013).

33. Gordon, R. et al. A simple magnetic separation method for high-yield isolation of pure primary microglia. Journal of neuroscience methods 194, 287–296, doi:10.1016/j.jneumeth.2010.11.001 (2011).

34. Turtzo, L. C. et al. Macrophagic and microglial responses after focal traumatic brain injury in the female rat. Journal of neuroinflammation 11, 82, doi:10.1186/1742-2094-11–82 (2014).

35. Trahanas, D. M., Cuda, C. M., Perlman, H. & Schwulst, S. J. Differential Activation of Infiltrating Monocyte-Derived Cells After Mild and Severe Traumatic Brain Injury. Shock 43, 255–260, doi:10.1097/SHK.0000000000000291 (2015).

36. Noristani, H. N. et al. RNA-Seq Analysis of Microglia Reveals Time-Dependent Activation of Specific Genetic Programs following Spinal Cord Injury. Front Mol Neurosci 10, 90, doi:10.3389/fnmol.2017.00090 (2017).

37. Matcovitch-Natan, O. et al. Microglia development follows a stepwise program to regulate brain homeostasis. Science 353, aad8670, doi:10.1126/science.aad8670 (2016).

38. Makinde, H. M., Cuda, C. M., Just, T. B., Perlman, H. R. & Schwulst, S. J. Nonclassical Monocytes Mediate Secondary Injury, Neurocognitive Outcome, and Neutrophil Infiltration after Traumatic Brain Injury. J Immunol 199, 3583–3591, doi:10.4049/jimmunol.1700896 (2017).

39. Schwarzmaier, S. M. & Plesnila, N. Contributions of the immune system to the pathophysiology of traumatic brain injury - evidence by intravital microscopy. Frontiers in cellular neuroscience 8, 358, doi:10.3389/fncel.2014.00358 (2014).

40. de Rivero Vaccari, J. P., Dietrich, W. D. & Keane, R. W. Activation and regulation of cellular inflammasomes: gaps in our knowledge for central nervous system injury. Journal of cerebral blood flow and metabolism : official journal of the International Society of Cerebral Blood Flow and Metabolism 34, 369–375, doi:10.1038/jcbfm.2013.227 (2014).

41. Edwards, P. et al. Final results of MRC CRASH, a randomised placebo-controlled trial of intravenous corticosteroid in adults with head injury-outcomes at 6 months. Lancet 365, 1957–1959, doi:10.1016/S0140-6736(05)66552-X (2005).

42. Roberts, I. et al. Effect of intravenous corticosteroids on death within 14 days in 10008 adults with clinically significant head injury (MRC CRASH trial): randomised placebo-controlled trial. Lancet 364, 1321–1328, doi:10.1016/S0140-6736(04)17188-2 (2004).

43. Wei, H. H. et al. NNZ-2566 treatment inhibits neuroinflammation and pro-inflammatory cytokine expression induced by experimental penetrating ballistic-like brain injury in rats. Journal of neuroinflammation 6, 19, doi:10.1186/1742-2094-6-19 (2009).

44. Cao, T., Thomas, T. C., Ziebell, J. M., Pauly, J. R. & Lifshitz, J. Morphological and genetic activation of microglia after diffuse traumatic brain injury in the rat. Neuroscience 225, 65–75, doi:10.1016/j.neuroscience.2012.08.058 (2012).

45. Krasemann, S. et al. The TREM2-APOE Pathway Drives the Transcriptional Phenotype of Dysfunctional Microglia in Neurodegenerative Diseases. Immunity 47, 566–581 e569, doi:10.1016/j.immuni.2017.08.008 (2017).

46. Boulanger, L. M. et al. Cellular and molecular characterization of a brain-enriched protein tyrosine phosphatase. The Journal of neuroscience : the official journal of the Society for Neuroscience 15, 1532–1544 (1995).

47. Lombroso, P. J., Naegele, J. R., Sharma, E. & Lerner, M. A protein tyrosine phosphatase expressed within dopaminoceptive neurons of the basal ganglia and related structures. The Journal of neuroscience : the official journal of the Society for Neuroscience 13, 3064–3074 (1993).

48. Bliss, T. V. & Collingridge, G. L. A synaptic model of memory: long-term potentiation in the hippocampus. Nature 361, 31–39, doi:10.1038/361031a0 (1993).

49. Silva, A. J. Molecular and cellular cognitive studies of the role of synaptic plasticity in memory. J Neurobiol 54, 224–237, doi:10.1002/neu.10169 (2003).

50. Braithwaite, S. P. et al. Regulation of NMDA receptor trafficking and function by striatal-enriched tyrosine phosphatase (STEP). The European journal of neuroscience 23, 2847–2856, doi:10.1111/j.1460-9568.2006.04837.x (2006).

51. Chin, J. et al. Fyn kinase induces synaptic and cognitive impairments in a transgenic mouse model of Alzheimer’s disease. The Journal of neuroscience : the official journal of the Society for Neuroscience 25, 9694–9703, doi:10.1523/JNEUROSCI.2980-05.2005 (2005).

52. Kurup, P. et al. Abeta-mediated NMDA receptor endocytosis in Alzheimer’s disease involves ubiquitination of the tyrosine phosphatase STEP61. The Journal of neuroscience : the official journal of the Society for Neuroscience 30, 5948–5957, doi:10.1523/JNEUROSCI.0157-10.2010 (2010).

53. Zhang, Y. et al. Genetic reduction of striatal-enriched tyrosine phosphatase (STEP) reverses cognitive and cellular deficits in an Alzheimer’s disease mouse model. Proceedings of the National Academy of Sciences of the United States of America 107, 19014–19019, doi:10.1073/pnas.1013543107 (2010).

54. Zhang, Y. et al. Reduced levels of the tyrosine phosphatase STEP block beta amyloid-mediated GluA1/GluA2 receptor internalization. Journal of neurochemistry 119, 664–672, doi:10.1111/j.1471-4159.2011.07450.x (2011).

55. Gold, J. M. & Harvey, P. D. Cognitive deficits in schizophrenia. Psychiatr Clin North Am 16, 295–312 (1993).

56. Bourne, C., Clayton, C., Murch, A. & Grant, J. Cognitive impairment and behavioural difficulties in patients with Huntington’s disease. Nurs Stand 20, 41–44, doi:10.7748/ns2006.05.20.35.41.c4146 (2006).

57. Anderson, C. A. & Arciniegas, D. B. Cognitive sequelae of hypoxic-ischemic brain injury: a review. NeuroRehabilitation 26, 47–63, doi:10.3233/NRE-2010-0535 (2010).

58. Ginhoux, F. et al. Fate mapping analysis reveals that adult microglia derive from primitive macrophages. Science 330, 841–845, doi:10.1126/science.1194637 (2010).

59. Schulz, C. et al. A lineage of myeloid cells independent of Myb and hematopoietic stem cells. Science 336, 86–90, doi:10.1126/science.1219179 (2012).

60. Gomez Perdiguero, E., Schulz, C. & Geissmann, F. Development and homeostasis of “resident” myeloid cells: the case of the microglia. Glia 61, 112–120, doi:10.1002/glia.22393 (2013).

61. Butovsky, O. et al. Identification of a unique TGF-beta-dependent molecular and functional signature in microglia. Nature neuroscience 17, 131–143, doi:10.1038/nn.3599 (2014).

62. Lavin, Y. et al. Tissue-resident macrophage enhancer landscapes are shaped by the local microenvironment. Cell 159, 1312–1326, doi:10.1016/j.cell.2014.11.018 (2014).

63. Ajami, B., Bennett, J. L., Krieger, C., Tetzlaff, W. & Rossi, F. M. Local self-renewal can sustain CNS microglia maintenance and function throughout adult life. Nature neuroscience 10, 1538–1543, doi:10.1038/nn2014 (2007).

64. Aguzzi, A., Barres, B. A. & Bennett, M. L. Microglia: scapegoat, saboteur, or something else? Science 339, 156–161, doi:10.1126/science.1227901 (2013).

65. Ajami, B., Bennett, J. L., Krieger, C., McNagny, K. M. & Rossi, F. M. Infiltrating monocytes trigger EAE progression, but do not contribute to the resident microglia pool. Nature neuroscience 14, 1142–1149, doi:10.1038/nn.2887 (2011).

66. Zeisel, A. et al. Brain structure. Cell types in the mouse cortex and hippocampus revealed by single-cell RNA-seq. Science 347, 1138–1142, doi:10.1126/science.aaa1934 (2015).

67. Goebel-Goody, S. M. et al. Genetic manipulation of STEP reverses behavioral abnormalities in a fragile X syndrome mouse model. Genes Brain Behav 11, 586–600, doi:10.1111/j.1601-183X.2012.00781.x (2012).

68. Poddar, R., Deb, I., Mukherjee, S. & Paul, S. NR2B-NMDA receptor mediated modulation of the tyrosine phosphatase STEP regulates glutamate induced neuronal cell death. Journal of neurochemistry 115, 1350–1362, doi:10.1111/j.1471-4159.2010.07035.x (2010).

69. Grabert, K. et al. Microglial brain region-dependent diversity and selective regional sensitivities to aging. Nat Neurosci 19, 504–516, doi:10.1038/nn.4222 (2016).

70. Dobin, A. et al. STAR: ultrafast universal RNA-seq aligner. Bioinformatics 29, 15–21, doi:10.1093/bioinformatics/bts635 (2013).

71. Eden, E., Navon, R., Steinfeld, I., Lipson, D. & Yakhini, Z. GOrilla: a tool for discovery and visualization of enriched GO terms in ranked gene lists. BMC Bioinformatics 10, 48, doi:10.1186/1471-2105-10-48 (2009).

